# Stabilization of fluctuating population dynamics via the evolution of dormancy

**DOI:** 10.1101/2024.09.12.612663

**Authors:** Zachary R. Miller, David Vasseur, Pincelli M. Hull

## Abstract

Dormancy is usually understood as a strategy for coping with extrinsically variable environments, but intrinsic population fluctuations also create conditions where dormancy is adaptive. By analyzing simple population models, we show that, very generally, population fluctuations favor the evolution of dormancy, but dormancy stabilizes population dynamics. This sets up a feedback loop that can enable the coexistence of alternative dormancy strategies. Over longer timescales, we show that evolution of dormancy to an evolutionary stable state can drive populations to the edge of stability, where dynamics are only weakly stabilized. We briefly consider how these conclusions are likely to apply in more complex community contexts. Our results suggest that chaos and high-amplitude population cycles are highly vulnerable to invasion and subsequent stabilization by dormancy, potentially explaining their rarity. At the same time, the propensity of ecological dynamics to fluctuate may be an underappreciated driver of the evolution of dormancy.

## Introduction

All organisms must cope with environments that vary in time. Physical conditions vary on many timescales, including daily, seasonal, and decadal cycles, and may be periodic or unpredictable (Huybers & Curry, 2006; Dillon *et al*., 2016; Park & Felmy, 2023). But in addition to abiotic forcings, organisms also face biotic conditions that can change dramatically through time. Fluctuations in prey availability, predation pressure, or competition intensity may amplify the effect of abiotic drivers or even arise in their absence (Ellner, 1985, 1987; Verin & Tellier, 2018; Munch et al., 2022; Ruf & Bieber, 2023). Because this kind of variability stems from population dynamics, strategies evolved in response to this variability not only shape the survival and fitness of individuals, but may potentially feed back to shape the character of the temporal variability itself.

Dormancy, a reversible suspension or reduction of metabolic activity, is perhaps the most widespread strategy used to contend with time-varying environments (Bewley, 1997; Wilsterman *et al*., 2021; Lennon *et al*., 2021). By entering dormancy, organisms reduce their metabolic demands and insulate themselves from unfavorable periods of high risk, such as drought, extreme temperatures, or low resource availability. Dormancy has clear adaptive value when unfavorable conditions are predictable, either because they occur regularly (e.g. diurnal cycles), set in slowly (e.g. resource depletion), or are preceded by reliable cues (e.g. seasonal changes) (Malik & Smith, 2006, 2008; Blath et al., 2021). In these cases, dormancy allows many organisms to substitute a period of no growth but low mortality for one where mortality risk would be high or where metabolic costs would outstrip fitness gains.

However, dormancy may provide a fitness benefit even in unpredictable environments as a bethedging strategy (Cohen, 1966; Kussell *et al*., 2005; Venable, 2007; Simons, 2011; Lennon *et al*., 2021; Blath et al., 2021). In this case, individuals enter and exit dormancy on a schedule – perhaps stochastic – that is independent of environmental conditions, with a chance of missing opportunities for growth and reproduction but also of avoiding periods of risk. By sacrificing potential fitness gains in good times to minimize fitness losses in bad times, such a strategy can reduce temporal variance in fitness, thereby increasing geometric mean fitness (long-term growth rate) in sufficiently variable and risky environments (Kussell & Leibler, 2005; Malik & Smith, 2008; Simons, 2011; Blath *et al*., 2021). A classic example comes from terrestrial plants, where many species produce seeds that exhibit heterogeneity in their germination timing, with some fraction remaining dormant through one or more germination opportunities (Cohen, 1966; Ellner, 1987, 1985). This has been predicted (Cohen, 1966; Bulmer, 1984; Ellner, 1985) and demonstrated (Clauss & Venable, 2000; Venable, 2007; Gremer & Venable, 2014) to function as an adaptive bet-hedging strategy against unpredictable risks such as drought.

There is an extensive theoretical literature on the adaptive value of dormancy as a bet-hedging strategy in different environmental contexts (Cohen, 1966; Bulmer, 1984; Ellner, 1985; Bär *et al*., 2002; Kussell et al., 2005; Kussell & Leibler, 2005; Malik & Smith, 2008; Blath et al., 2021; Măgălie *et al*., 2023). However, most of this theory is concerned with hedging against extrinsic sources of variation. While it is well-understood that density-dependent population dynamics can may impact the evolution of dormancy in these contexts (Ellner, 1985; Gremer & Venable, 2014; Kortessis & Chesson, 2019), we know little about how organisms use dormancy to cope with fluctuations arising from population or community dynamics themselves, and what the consequences of such strategies might be.

Yet, intrinsic population fluctuations may represent an important source of variability for many organisms. Theoretical models of ecological dynamics suggest that population cycles and chaos – bounded, aperiodic dynamics that depend sensitively on initial conditions – should be common (Hastings *et al*., 1993; Munch et al., 2022; Robey et al., 2024). Stable predator-prey cycles were one of the first theoretical predictions in ecology (Lotka, 1920; Volterra, 1926), and cycles and chaos are typical dynamics in food chain models (Gilpin, 1979; Hastings et al., 1993; Vandermeer, 1993; Gross *et al*., 2005). Endogenous fluctuations also occur in a large class of discrete time population models with strong (overcompensatory) density dependence (Geritz & Kisdi, 2004; Devaney, 2019), and may be a very general feature of community models with many species and strong interactions (Ispolatov *et al*., 2015; Roy et al., 2020). These mechanisms give rise to dynamics that fluctuate without abiotic forcing. From the perspective of individual organisms, they generate conditions where growth and mortality rates vary, potentially dramatically and unpredictably.

Cycles and chaos have been demonstrated in experimental populations and simplified communities (Ellner & Turchin, 1995; Turchin & Ellner, 2000; McCauley et al., 1999; Benincà et al., 2008; Blasius *et al*., 2020; Munch et al., 2022; Robey et al., 2024), but ecologists have long noted a gap between theoretical models, where cycles and chaos appear to be “the rule rather than the exception” (Hastings *et al*., 1993), and natural communities, where evidence for these dynamics, especially chaos, is limited (Ellner & Turchin, 1995; Sibly *et al*., 2007; Robey *et al*., 2024). This mismatch fuels long-standing debate over their ecological reality and relevance (Berryman & Millstein, 1989; Munch et al., 2022). More recently, careful analyses of extensive time-series data have suggested that chaos is not rare, but may be generally weak or intermittent, with many populations poised near the “edge of chaos” (Turchin & Ellner, 2000; Munch et al., 2022; Rogers et al., 2022, 2023). This combination of extensive evidence for the potential for fluctuating population dynamics, together with evidence that these dynamics are perhaps rare and rarely strong, has spurred interest in mechanisms that might stabilize or suppress fluctuations (Berryman & Millstein, 1989; Doebeli, 1997; Ebenman et al., 1997). Dormancy has repeatedly been proposed as such a mechanism (Ruxton & Rohani, 1998; McCauley *et al*., 1999; Hadeler, 2008; Kuwamura *et al*., 2009; Tan *et al*., 2020), because, as a bet-hedging strategy, it acts precisely to dampen variation in growth rates.

In short, fluctuating population dynamics, which are widely predicted by theory, create conditions where dormancy is highly adaptive. At the same time, widespread dormancy is expected to weaken or eliminate these dynamics, consistent with most observations in nature.

These two predictions suggest an inevitable feedback loop between population fluctuations and dormancy. This possibility was noted by Lalonde & Roitberg (2006), but the resulting dynamics have been scarcely studied, and only in relatively complex models that are challenging to analyze (Lalonde & Roitberg, 2006; Wen et al., 2022).

In this Letter, we explore the potential feedbacks between population fluctuations and the evolution of dormancy as a bet-hedging strategy. Using a highly simplified model of fluctuating population dynamics to illustrate our results, we show that (i) dormancy has a stabilizing effect on dynamics, (ii) dormancy is an adaptive strategy that has positive invasion fitness when dynamics fluctuate, (iii) feedbacks between different dormancy strategies can enable their coexistence, and (iv) the evolution of dormancy can drive population dynamics to the edge of stability. We show that each of these four conclusions applies quite generally to discrete time population models, and likely also to community models where fluctuations arise from interspecific interactions. Our results indicate that, given the tendency of population dynamics to fluctuate, these dynamics may be an underappreciated driver of the evolution and diversification of dormancy strategies. And given that dormancy appears readily evolvable across the tree of life, we suggest that this simple strategy may provide one general explanation for the rarity of high amplitude population cycles and strong chaos in nature.

## Model

As a minimal model for the interaction of population fluctuations and dormancy, we consider the discrete-time dynamics

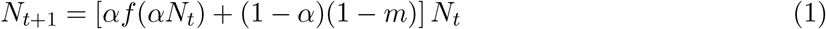

describing a species with non-overlapping generations (Fig. 1). In this model, *N*_*t*_ is the total population size at time *t*; *α* is the fraction of individuals that are active (not dormant) in each generation; *f* is the per capita net growth rate of active individuals, which depends on the number of active individuals at time *t*; and *m* is the per generation mortality risk for dormant individuals. Eq. 1 naturally models a life cycle where all individuals pass through a seed bank stage, and some fraction of individuals remain there in a dormant state in each generation (Cohen, 1966; Ellner, 1987). We assume that this fraction, 1 − *α*, is constant over time, and does not depend on any environmental conditions or other cues. At the individual level, transitions out of dormancy can be viewed as stochastic; effectively, we assume that each individual remains dormant with probability 1 − *α* in each generation and emerges from dormancy with probability *α*, as depicted in Fig. 1. Consequently, there is a persistent pool of dormant individuals across generations, of size (1 − *α*)*N*_*t*_, but the time any individual spends dormant is a geometrically-distributed random variable with mean 1*/α*.

**Figure 1:**
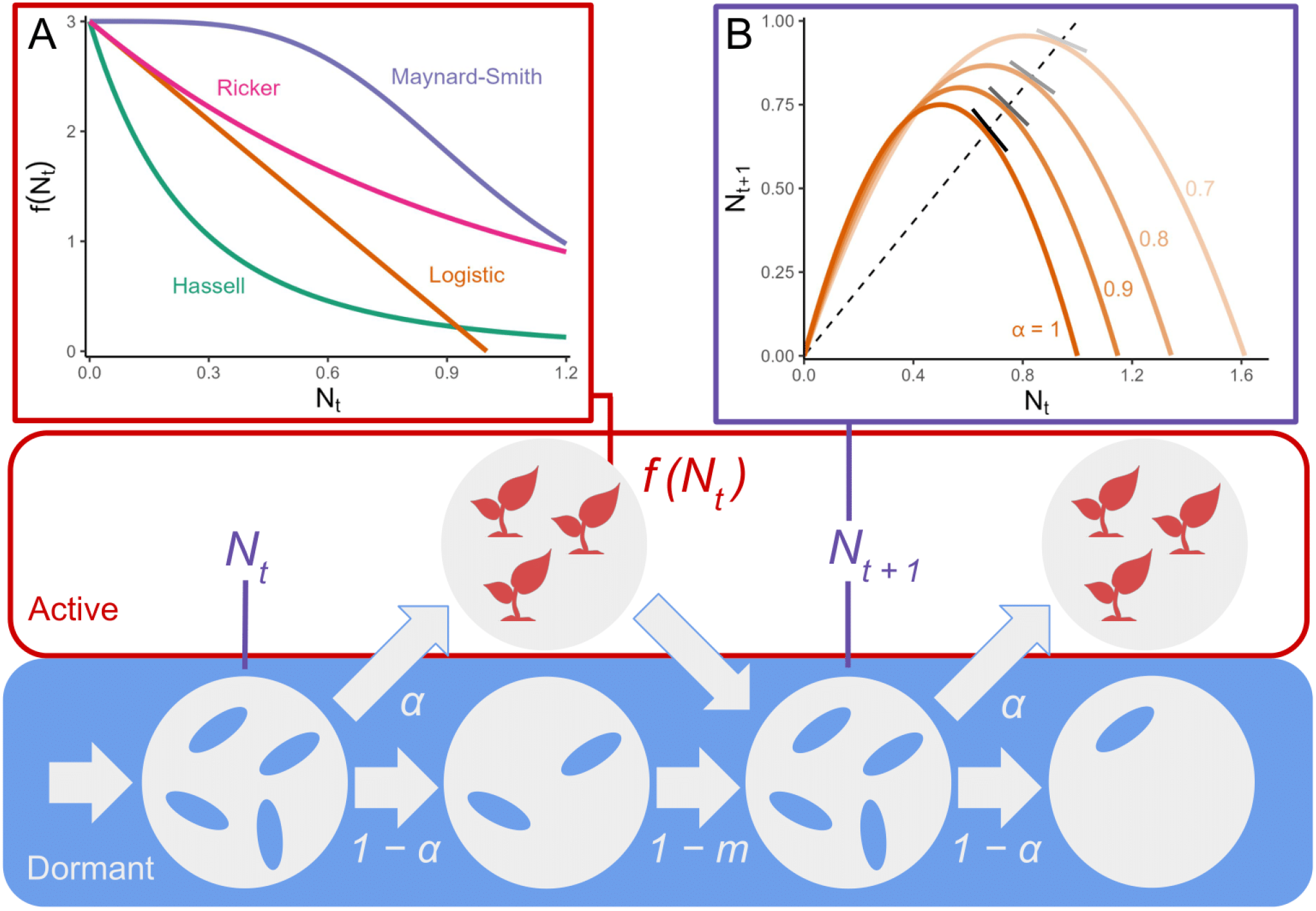
Graphical model of Eq. 1. In each generation (left to right), a fraction *α* of the population becomes active (red), and a fraction 1 − *α* is dormant (blue). Active individuals have a per capita growth rate that depends on the total active population size. The form of this dependence is specified by a function *f* ; several choices of *f* corresponding to well-known population models are shown (Inset A). In Eq. 1, the entire population (combining dormant and active fractions) is censused at the end of each generation (purple). The contribution of both active and dormant fractions results in a map between the total population size at times *t* and *t* + 1. Different values of *α* produce a family of maps, illustrated for the logistic model (Eq. 2) in Inset B. The population possesses an equilibrium where *N*_*t*+1_ = *N*_*t*_ (dashed line); the position and dynamical stability of this equilibrium depends on the value of *α*. In general, as *α* decreases (increasing dormancy), the slope of *N*_*t*+1_ as a function of *N*_*t*_ becomes shallower at the equilibrium point, corresponding to more stable dynamics.

While this model of dormancy is highly simplified, similar assumptions are often used to describe dormancy in organisms such as annual plants (Cohen, 1966; Bulmer, 1984; Ellner, 1985, 1987; Gremer & Venable, 2014) and bacteria (Kussell *et al*., 2005; Kussell & Leibler, 2005; Lennon *et al*., 2021), which can exhibit stochastic dormancy strategies (Balaban *et al*., 2004; Buerger et al., 2012). When *α* = 1, we recover a standard model of density-dependent population dynamics, with no individuals dormant across generations. For convenience, we call this the “no dormancy” case, although we note that all individuals initially pass through the seed-bank stage (e.g. as an overwintering stage).

To reflect negative density dependence due to competition or crowding, we assume that the net growth rate *f* is a decreasing function of the (active) population size. If density dependence is sufficiently strong, the model may be overcompensating, meaning that *N*_*t*+1_ has a maximum as a function of *N*_*t*_. Overcompensating models can exhibit complex non-equilibrium population dynamics, including cycles or chaos (May & Oster, 1976; Geritz & Kisdi, 2004; Devaney, 2019).

With no dormancy (*α* = 1), different choices of *f* yield well-known population models, such as the Hassell (Hassell, 1975), Ricker (Ricker, 1954), and Maynard-Smith models (Smith & Slatkin, 1973) (Fig. 1A). For each of these models, the density-dependent growth rate takes the form *f*_*r*_(*N*) = *rg*(*N*), where *r >* 0 is typically interpreted as a maximum or intrinsic growth rate. We focus throughout on models of this form, which we call “separable” models, although our conclusions likely apply more generally (see SI Section A). If the dynamics are overcompensating – in all of these cases, for example – increasing *r* can produce a bifurcation, where the long-term dynamics shift from a stable equilibrium to cyclic or chaotic fluctuations around an unstable equilibrium (Geritz & Kisdi, 2004). The bifurcation occurs at a point *r*_*c*_ where the slope of *N*_*t*+1_ as a function of *N*_*t*_, evaluated at an equilibrium abundance 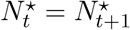, exceeds 1 in absolute value (Fig. 1B).

As a concrete example of a model with separable and overcompensating dynamics, we will consider the discrete logistic model with dormancy

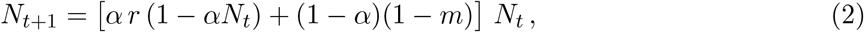

which corresponds to choosing *f*_*r*_(*N*) = *r*(1 − *N*) in Eq. 1. Here, we are assuming that *N* is measured in units of carrying capacity for the (active) population. The discrete logistic model with no dormancy provides one of the simplest, oldest, and best-studied examples of chaotic dynamics in ecology (May & Oster, 1976; Hastings et al., 1993; Robey et al., 2024). Moreover, the logistic model typifies the dynamical properties of other overcompensating models (Devaney, 2019). For *r* > *r*_*c*_ = 3 the dynamics without dormancy become cyclic – with increasing period and amplitude, following a period-doubling cascade (May & Oster, 1976; Devaney, 2019) – and eventually chaotic. Above *r* = 4, the fluctuations become large enough to drive the population extinct.

## Results

### Dormancy stabilizes fluctuating dynamics

In overcompensating models, high intrinsic growth rates produce fluctuating dynamics as populations begin to over- and under-shoot their carrying capacity. Dormancy, by reducing temporal variation in growth rates, can temper such booms and busts.

In fact, dormancy can qualitatively stabilize these dynamics. We show (SI Section A) that if mortality risk in dormancy (*m*) is sufficiently low, then decreasing *α* in Eq. 1 raises the bifurcation point *r*_*c*_. That is, allowing some fraction of individuals to go dormant between generations increases the range of intrinsic growth rates that permit a stable equilibrium.

For the logistic model with dormancy (Eq. 2), we can precisely map out this stabilizing effect. By collecting density-independent terms in Eq. 2 and re-scaling the population size (see SI Section A.3), we obtain an effective intrinsic growth rate for the population, which characterizes its dynamics:

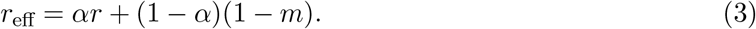

The effective growth rate, *r*_eff_, reflects the intrinsic growth rate of the active fraction of the population (first term) and the survival probability of the dormant fraction (second term). The behavior of Eq. 2 with a certain *r*_eff_ is qualitatively equivalent to a logistic model with no dormancy (*α* = 1) and *r* = *r*_eff_. Because the behavior of the logistic model without dormancy is well-characterized for all intrinsic growth rates (Devaney, 2019), we can use *r*_eff_ to determine the dynamics of Eq. 2 for any *α, r*, and *m*. For example, Fig. 2, shows the dynamics that result for varying *α* and *r* (for a fixed *m*).

**Figure 2:**
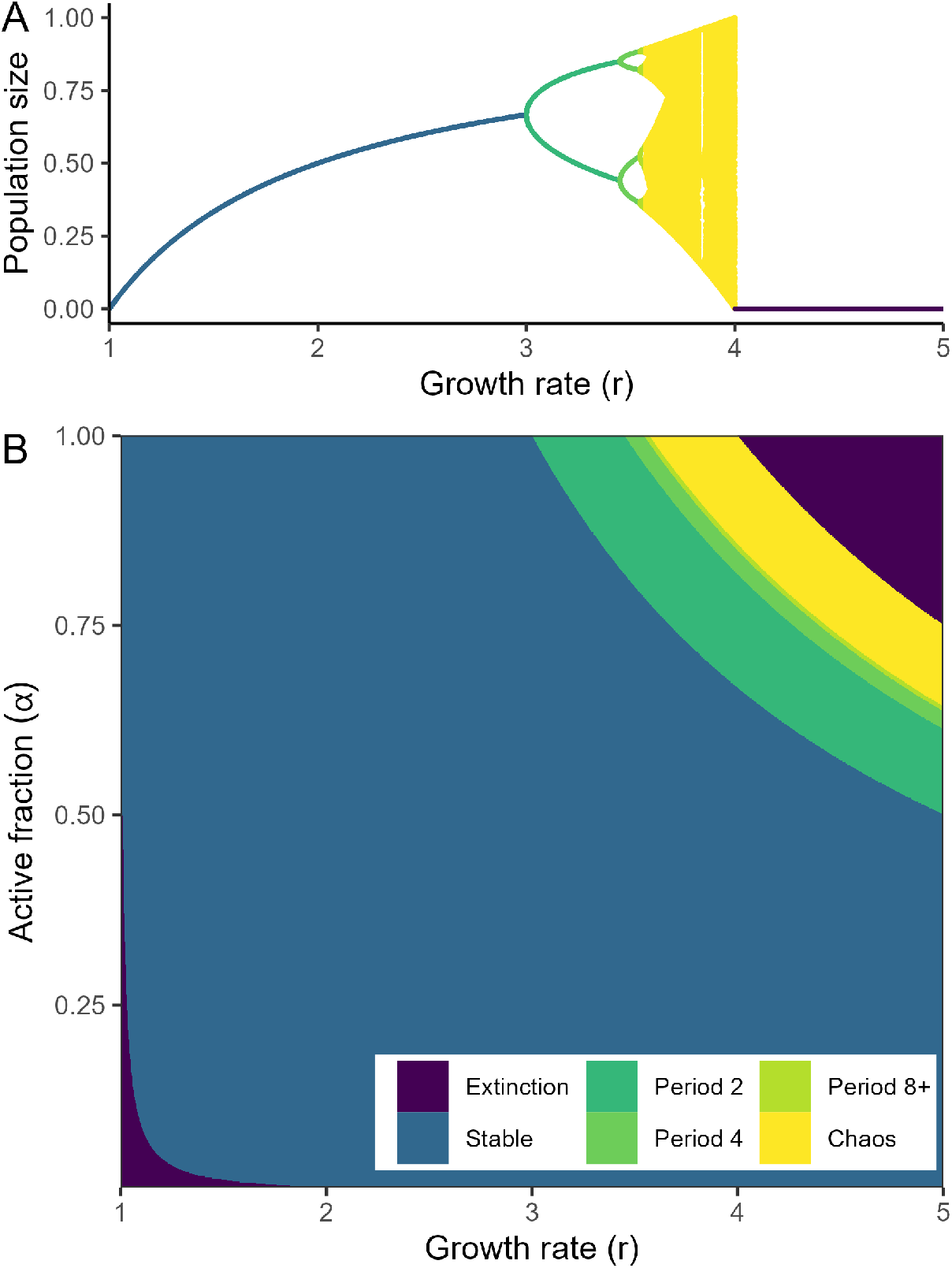
Bifurcation diagrams for the logistic model (Eq. 2). (A) Steady state population size(s) are plotted against *r* for the standard logistic model with *α* = 1. For *r <* 3 the model possesses a stable equilibrium. At *r* = 3 the dynamics undergo a bifurcation and stable oscillations (green) emerge. Above *r* ≈ 3.57, the dynamics are chaotic (yellow), and above *r* = 4, the population fluctuates to extinction. (B) Qualitative steady state dynamics for different combinations of *r* and *α* (with *m* = 0.01). Dormancy increases from top to bottom. Each point in (B) can be mapped to a point in (A) following Eq. 3. For lower values of *α* (more dormancy), population fluctuations arise at higher growth rates (*r*). Note that for very low *α* and *r*, the effective population growth rate falls below 1, leading to extinction.

Fig. 2 illustrates that as individuals spend more time in dormancy (decreasing *α*), the population dynamics remain stable for larger intrinsic growth rates. Indeed, by solving Eq. 3 for the bifurcation point in the baseline model (*r*_*c*_ = 3), the bifurcation in Eq. 2 occurs when *r* surpasses

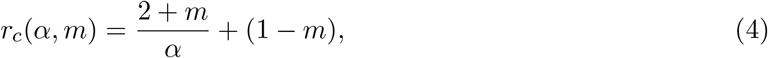

which is a decreasing function of *α*. Similarly, the onset of chaos (yellow) and of fluctuations to extinction (violet) are always postponed by increasing dormancy.

### Dormancy invades fluctuating dynamics when rare

In the presence of fluctuating dynamics, dormancy is an adaptive strategy. To establish this, we calculate the invasion growth rate (IGR) (Metz *et al*., 1992; Grainger et al., 2019) for a rare ecotype with an active fraction *α*^′^ (“high dormancy”) in a fluctuating resident population with no or lower dormancy (1 ≥ *α* > *α*^′^). We assume the resident population size, *N*_1_, *N*_2_, …, has reached a stationary state (limit cycle or chaotic attractor) where the per capita growth rate of the resident has a geometric mean of one over the long term. Equivalently, defining *h*_*t*_ = *αf*_*r*_(*αN*_*t*_) + (1 − *α*)(1 − *m*), we have that 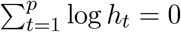, where *p* is the period of the resident population fluctuations (for chaotic dynamics, we take *p* → ∞). In this setting, the log long-term growth rate (Metz *et al*., 1992) for a rare ecotype with the same *r* and *m* but *α*^′^ < *α* is

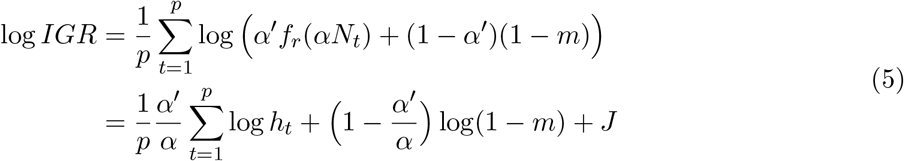

(see SI Section B for a detailed derivation). The log IGR can be understood as a sum of three components (bottom line): a term proportional to the resident’s mean growth rate, which is zero because the resident population is stationary; a term 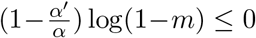, quantifying the mortality cost of increasing dormancy; and a term *J* > 0, quantifying the bet-hedging benefit of increasing dormancy. The term *J* is generally unwieldy, but, crucially, this benefit term is strictly positive as a consequence of nonlinear averaging over times when the population booms and busts (Ruel *et al*., 1999). Intuitively, *J* quantifies the fact that reduced growth during population booms is more than offset by reduced losses during busts. Thus, the IGR reflects a balance of the costs and benefits of dormancy. If mortality risk in dormancy, *m*, is sufficiently small, then the benefit *J* exceeds the mortality cost and log *IGR* > 0.

This means that a small subpopulation with increased dormancy can grow when rare in a fluctuating resident population. To reach this conclusion, we assumed that the resident and invader have equal intrinsic growth rates and that mortality in dormancy is low. However, both assumptions can be relaxed, to an extent that depends on the underlying population model. In general, larger fluctuations generate a greater bet-hedging benefit for dormancy (larger *J*). Sufficiently large fluctuations can favor invasion even when the invader has a lower intrinsic growth rate than the resident, and despite moderate mortality risk in dormancy. For example, Fig. 3 illustrates a scenario where an invader with *r*^′^ = 3.8 and *m* = 0.05 grows rapidly in a chaotic resident population with *r* = 3.9 and no dormancy. In the SI (Fig. S1), we numerically calculate the *IGR* in the logistic model for an invader with varying *α*^′^ and *m* in a resident population with no dormancy, for several values of *r*. Overall, as *r* increases, the resident population fluctuations increase in amplitude, and high-dormancy invaders are favored despite considerable mortality in dormancy. However, with non-negligible risk in dormancy, there is a lower limit on *α*^′^, below which dormancy is disfavored. Moderate dormancy generates a bet-hedging advantage that outweighs the mortality cost, but decreasing *α*^′^ eventually leads to increasing exposure to mortality with diminishing returns from bet-hedging.

**Figure 3:**
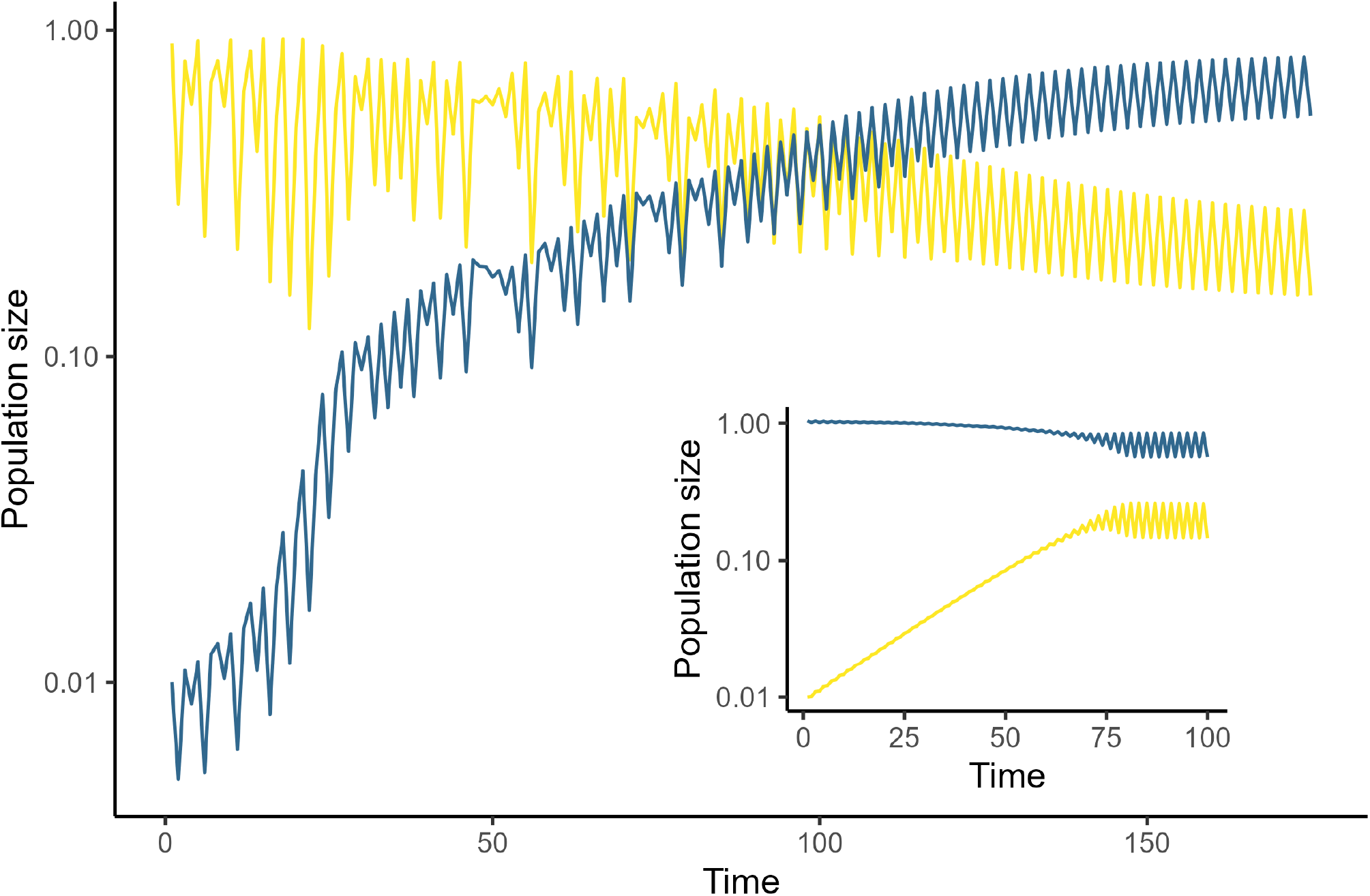
Mutual invasibility between low- and high-dormany ecotypes in the logistic model (Eq. 2). Main panel: Dynamics of a resident population (yellow) with parameters *r* = 3.9 and *α* = 0 invaded by a rare ecotype (blue) with *r*^′^ = 3.8, *m* = 0.05, and *α*^′^ = 0.7. The resident dynamics are chaotic with high-amplitude fluctuations; this generates a positive IGR for the high-dormancy ecotype. Once the invader becomes sufficiently abundant (around *t* = 100), the dynamics of both ecotypes shift from chaos to period-two oscillations with a low amplitude. Inset: Invasion of the high-dormancy ecotype (blue) by the low-dormancy type (yellow). The high-dormancy ecotype approaches a stable equilibrium in isolation, but the invasion of the low-dormancy ecotype induces oscillations. Because both ecotypes can invade when rare, they coexist robustly. Note the log scale of the y-axis in both panels.

### Dormancy enables coexistence via negative feedback

Our *IGR* calculation reveals an advantage for dormancy when rare. But as a subpopulation with higher dormancy grows, we have seen that it will begin to stabilize the population dynamics, potentially even suppressing fluctuations entirely. What happens as the proliferation of dormancy suppresses the temporal variability that underpins its growth rate advantage?

To answer this question, we consider the IGR of the original resident population, assuming it eventually becomes rare compared to the invader. We assume again that the original resident has an active fraction *α* (“low dormancy”) and the invader has *α*^′^ *< α* (“high dormancy”), with *r* and *m* equal for both types. Whether the original resident type can reciprocally invade depends on the intrinsic population dynamics of the high-dormancy ecotype. If *α*^′^ is low enough to fully stabilize the dynamics, then, in isolation, this population would reach an equilibrium size *N*^⋆^ satisfying *α*^′^*f*_*r*_(*α*^′^*N*^⋆^) + (1 − *α*^′^)(1 − *m*) = 1. At this equilibrium point, the *IGR* for the low-dormancy type is given by *IGR* = *αf*_*r*_(*α*^′^*N*^⋆^) + (1 − *α*)(1 − *m*). Subtracting the former equation from the latter, we have *IGR*−1 = (*α−α*^′^)(*f*_*r*_(*α*^′^*N*^⋆^)+*m*), which implies that *IGR* > 1 whenever *α* > *α*^′^. Consequently, if the high-dormancy population exhibits a stable equilibrium in isolation, then the original resident with lower dormancy can also invade when rare, due to a growth advantage in constant environments (Fig. 3b).

In this case, there is mutual invasibility between the two strategies (Grainger *et al*., 2019). The low-dormancy ecotype generates fluctuations that favor the high-dormancy type, and the high-dormancy type creates constant conditions that favor lower dormancy. This negative frequency-dependent feedback allows stable coexistence of the two strategies (Fig. 3), despite the fact that there are otherwise no niche differences between them (i.e., we assume that each active individual of either ecotype decreases the growth rate of both equally).

If fluctuations persist even when the high-dormancy type dominates the population, the situation is more complicated. Coexistence may still be possible, or the high-dormancy type may exclude the low-dormancy strategy. The outcome depends again on whether the IGR for low dormancy is greater or less than one, but calculating the IGR requires the distribution of population sizes for the fluctuating high-dormancy population. This distribution depends on the form of the model and may be challenging to calculate. Generally, however, as dormancy suppresses population fluctuations more strongly (i.e. smaller amplitude fluctuations), an advantage accrues to the low-dormancy strategy when rare, favoring coexistence.

### Dormancy drives evolution to the edge of stability

We have so far examined the interaction between two fixed dormancy strategies. A natural last question is how the level of dormancy will evolve over the long term in a population prone to fluctuating dynamics. Once again, we fix *r* and *m*, with *r* large enough that the population fluctuates with no dormancy and *m* small enough that dormancy can invade. We assume that *α* can evolve through the invasion of mutant ecotypes with varying dormancy strategies. If invasions are sufficiently rare so that the model dynamics always reach a stationary state between invasions, then the long-term dynamics of the dormancy trait value *α* are determined by the geometry of a pairwise invasibility plot (PIP), indicating which mutant trait values can invade for any resident value (Metz *et al*., 1992; McGill & Brown, 2007). The PIPs in Fig. 4 show that for the logistic model (Eq. 2), these dynamics possess a unique evolutionary stable strategy (ESS) – a value *α*^⋆^ that can resist invasion by any other dormancy strategy (vertical dashed lines). Furthermore, the ESS value *α*^⋆^ invades adjacent trait values (horizontal dashed lines), indicating that the ESS can be reached via gradual evolution. The same qualitative features are shared by other overcompensating models (SI Section D).

**Figure 4:**
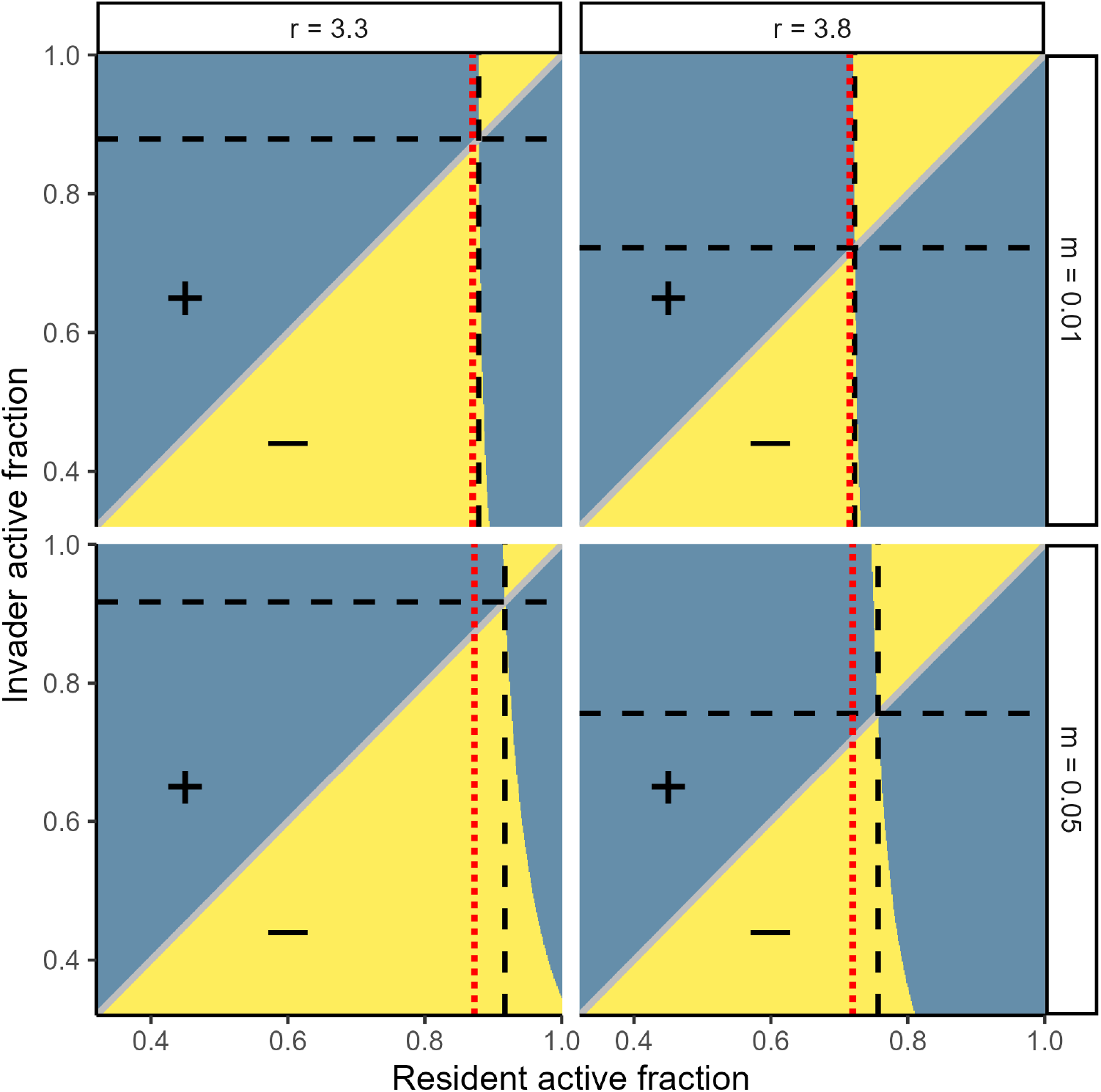
Pairwise invasibility plots for the trait value *α* in the logistic model (Eq. 2), with different fixed values of *r* and *m*. Blue regions indicate that the active fraction *α*^′^ on the y-axis can invade the resident active fraction *α* on the x-axis. Yellow regions indicate that the resident can resist invasion. For each combination of *r* and *m*, there is a unique value *α*^⋆^ that is not invasible by any other *α* value (vertical dashed line), and that can invade every other value (horizontal dashed line). These are evolutionary stable strategies (ESS). As discussed in the main text, the ESS typically sits close to – but just above – the value *α*_*c*_ where the dynamics begin to fluctuate (red dotted lines). For small values of *m* (e.g. top row), *α*^⋆^ ≈ *α*_*c*_.

When there is low mortality risk in dormancy (*m* ≪ 1), the ESS value *α*^⋆^ lies very close to a bifurcation point in *α*. Low values of *α*, corresponding to high dormancy, stabilize the dynamics. As *α* increases, there is necessarily a bifurcation point *α*_*c*_ where stability is lost (as we assume that increasing *α* all the way to one produces fluctuating dynamics). Populations with *α* > *α*_*c*_ fluctuate, and therefore can be invaded by mutants with lower *α* (more dormancy). Conversely, populations with *α* < *α*_*c*_ are stable; in this case, we have shown that there is a growth rate advantage for mutants with higher *α* (less dormancy), which have a higher effective growth rate. Only *α*_*c*_ is noninvasible from above and below. With non-negligible mortality in dormancy, *α*^⋆^ is generically slightly higher than *α*_*c*_, because the bet-hedging benefit of dormancy only outweighs the mortality cost once resident population fluctuations grow sufficiently large.

A population with *α* ≈ *α*_*c*_ is poised at the boundary between stable and unstable dynamics. At this bifurcation point, the dynamics approach an equilibrium that is only weakly stabilized. Formally, the dynamics at this point have a Lyapunov exponent of zero, indicating that trajectories converge to the equilibrium very slowly. Positive Lyapunov exponents imply chaotic dynamics, so systems with a Lyapunov exponent near zero are situated at the “edge of chaos” or “edge of stability” (Hastings *et al*., 1993; Turchin & Ellner, 2000; Munch *et al*., 2022). More generally, for *m* > 0, we expect the ESS to be slightly above the bifurcation point. This means that populations at the ESS will typically exhibit low amplitude fluctuations around an unstable equilibrium, with a small but negative Lyapunov exponent.

#### Box 1

Fluctuating population dynamics also occur in continuous-time models of multiple interacting species, especially when species interactions are strong (Gross *et al*., 2005; Ispolatov *et al*., 2015; Robey *et al*., 2024). Our conclusions for discrete-time population models in the main text apply broadly to continuous-time multispecies models, as well.

As a simple model of stochastic dormancy in continuous time, analogous to Eq. 1, we consider the dynamics of *S* species given by

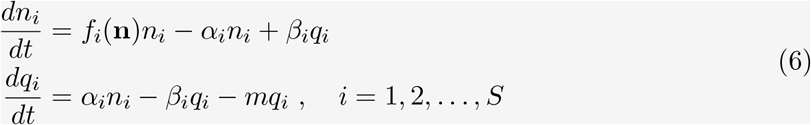

where *n*_*i*_ is the active population size of species *i, q*_*i*_ is its dormant population size, and *f*_*i*_ gives the per capita growth rate of species *i* as a function of **n** = (*n*_1_, *n*_2_, …, *n*_*S*_). Individuals of each species transition in and out of dormancy at rates *α*_*i*_ and *β*_*i*_, respectively, and *m* is the mortality rate in the dormant state. Like Eq. 1, the model in Eq. 6 describes a system where individuals enter and exit a dormant state stochastically and independently of one another or any cues (Hadeler, 2008; Hadeler et al., 2009).

We briefly consider how our main results apply in this setting.

**Invasion**

As in discrete time, population fluctuations in continuous time confer a growth rate advantage to dormancy. Consider a resident community with no dormancy (i.e. *α*_*i*_ = *β*_*i*_ = 0 for all *i*) that fluctuates in a limit cycle or chaotic attractor. In the SI (Section C), we calculate the IGR for a rare “mutant” of any species *j*, which shares the growth rate function *f*_*j*_ but has a dormant state 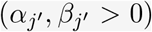. We show that regardless of the specific community dynamics, if the mortality rate *m* is sufficiently low, then this IGR is always positive in two limits that bookend all possible scenarios: 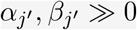 (very frequent transitions in and out of dormancy) and 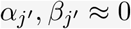 (very infrequent transitions).

**Stabilization**

The effects of dormancy on stability are more complex here, because different species may have different dormancy transition rates (Hadeler, 2008). In a simple scenario where *α*_*i*_ = *α* and *β*_*i*_ = *β* across all species, it is known that dormancy is invariably sta-bilizing – if the underlying dynamics 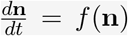 fluctuate around an unstable equilibrium, then introducing dormancy dampens high-frequency oscillations and can even qualitatively stabilize the equilibrium (Hadeler, 2008; Hadeler et al., 2009). More generally, dormancy can potentially stabilize or destabilize dynamics, depending on the specific model (Hadeler, 2008; Hadeler *et al*., 2009). However, we hypothesize that when strongly-interacting members of the community have a dormant state, this will often have a stabilizing effect by dampening oscillatory dynamics (e.g. Fig. 5).

**Figure 5:**
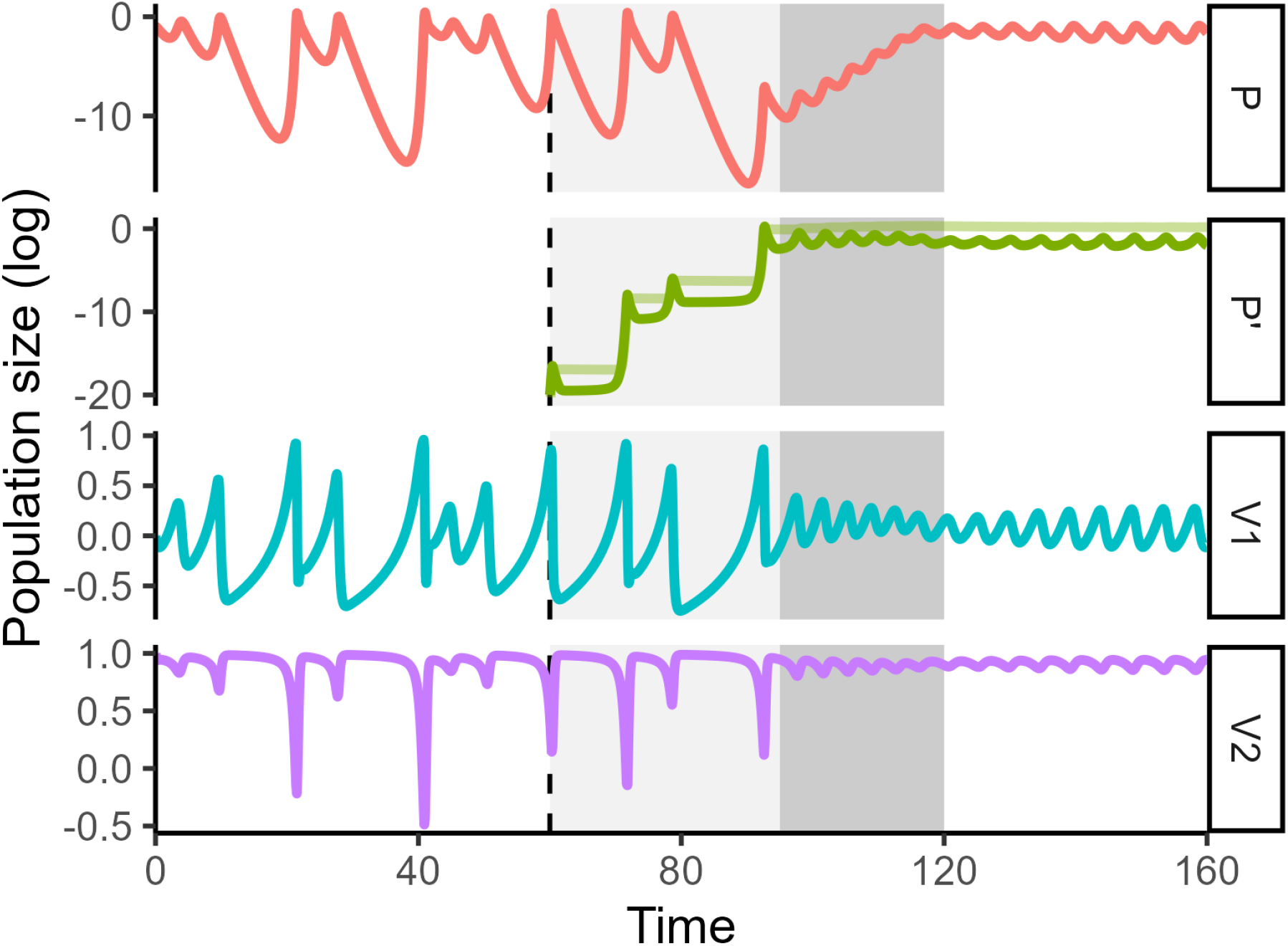
Chaotic food web dynamics (Eq. 7) invaded by a predator with dormancy. At time *t* = 60, a “mutant” predator (*P* ^′^, green) invades the system at very low abundance. For *P* ^′^, the active population 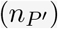 is indicated by a dark line, and the dormant population 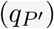 by a light line. Parameter values are *r*_1_ = *r*_2_ = *r*_3_ = 10, *a*_11_ = *a*_12_ = *a*_22_ = *a*_21_ = 1, *a*_13_ = 10, *a*_21_ = 1.5, *a*_31_ = 5, *a*_32_ = 0.5, and *α* = 1.5, *β* = 0.015, *m* = 0.01 for the invader *P* ^′^.

**Coexistence**

If the invasion of a mutant with dormancy stabilizes the dynamics, coexistence between the mutant and resident ecotypes may be possible due to the same feedback identified in the main text: The mutant with dormancy experiences a positive IGR when dynamics fluctuate, but once this ecotype becomes abundant, fluctations are suppressed, returning an advantage to the ecotype without dormancy. Assuming *m >* 0, the no-dormancy ecotype necessarily has a positive IGR if the community dynamics approach an equilibrium after the invader with dormancy establishes. Under these conditions, the two ecotypes will coexist, typically in a low amplitude limit cycle.

**Long-term evolution**

The dynamics above set the stage for the trait values that control dormancy (*α*_*i*_ and *β*_*i*_) to evolve toward a bifurcation point (“the edge of chaos”). Fully characterizing this eco-evolutionary feedback requires greater understanding of the stabilizing effects of dormancy in a community context, making this a compelling goal for future research.

We illustrate these results in a simple food web model with chaotic dynamics. The following Lotka-Volterra dynamics studied by Gilpin (1979) model a predator species (*P*) that consumes two competing prey (*V*_1_ and *V*_2_):

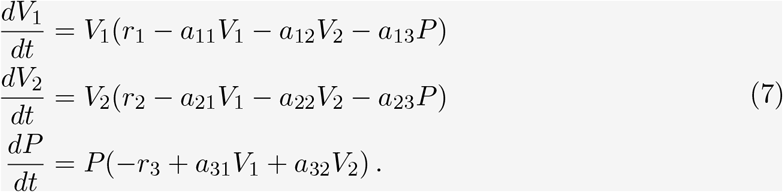

For parameters shown in Fig. 5, the community dynamics are chaotic. At time *t* = 60, we introduce a rare species *P* ^′^ that is identical to the predator, but has a dormant state. Fig. 5 shows that *P* ^′^ rapidly rises in abundance (light gray window) owing to a positive IGR. Once the invader becomes more abundant than the resident predator, the community dynamics begin to stabilize, exhibiting reduced booms and busts (dark gray window). This reduces the invader’s growth rate advantage over the resident predator, which rebounds in abundance. Eventually, the two ecotypes *P* and *P* ^′^ coexist as the entire community settles into weak oscillations.

## Discussion

Dormancy is a widespread strategy for coping with fluctuating conditions, whether extrinsically or intrinsically driven (Wilsterman *et al*., 2021; Lennon et al., 2021). Intrinsic population fluctuations are a typical prediction of ecological models (Hastings *et al*., 1993; Munch *et al*., 2022). Here, we have shown that even a simplistic, stochastic dormancy strategy can be highly adaptive under such dynamics. We have also shown that the invasion of dormancy into a fluctuating population has a stabilizing effect. The resulting feedback between a high-dormancy ecotype, which benefits from fluctuations but stabilizes them, and a low-dormancy ecotype, which destabilizes dynamics but benefits from stability, can allow stable coexistence of different strategies, and can ultimately favor trait evolution toward the edge of stability.

For clarity and mathematical tractability, we focused our analysis on a class of discrete-time population models. However, in Box 1, we explore how our four main conclusions apply to multispecies dynamics in continuous time (i.e. for species that reproduce continually with overlapping generations). Conceptually, we expect our main conclusions to be robust to the specific form of dynamics: Because fitness fluctuations generically reduce a population’s long-term (multiplicative) growth rate, fluctuating population dynamics should generally favor the evolution of dormancy. And because dormancy makes a population’s dynamics less sensitive to per capita fitness fluctuations, dormancy should generally tend to stabilize population dynamics. However, characterizing precisely when and how dormancy can stabilize more complex community dynamics remains an open problem (Hadeler, 2008; Hadeler et al., 2009; Wen et al., 2022).

We also considered a highly simplified model of dormancy, where activity is a binary state (active or dormant) and transitions in and out of dormancy occur at a constant probability through time, with no dependence on population size or other factors. In reality, most organisms employ more complex dormancy strategies, ramping metabolic activity up or down in response to environmental cues (Bär *et al*., 2002; Kussell & Leibler, 2005; Malik & Smith, 2006). For example, many species can detect early indicators of resource scarcity and enter dormancy to avoid starvation. While these kinds of adaptive behaviors are not included in our models, they would most likely only increase the benefit of dormancy under population fluctuations, compared to a pure bet-hedging strategy (see, e.g., SI Section C.2). We also hypothesize that more sophisticated dormancy strategies are even more strongly stabilizing, by reducing or eliminating large population crashes.

Interest in dormancy is growing as biologists increasingly recognize its ubiquity and its importance for understanding pressing biological problems, from climate resilience to antibiotic resistance (Balaban *et al*., 2004; Kussell *et al*., 2005; Lennon *et al*., 2021; Wilsterman *et al*., 2021; Wisnoski & Lennon, 2021). Yet, even as our knowledge of the physiological mechanisms and individual-level costs and benefits of dormancy has expanded, we have only just begun to explore the population- and community-level consequences of dormancy. While elements of the eco-evolutionary feedback we identify have been recognized before (e.g. the adaptive value of dormancy in fluctuating populations (Ellner, 1987; Ebenman *et al*., 1997; Malik & Smith, 2008; Tan *et al*., 2020; Blath *et al*., 2021) and the stabilizing effects of dormancy (Hadeler, 2008; Hadeler et al., 2009; Kuwamura et al., 2009; Wen *et al*., 2022)), the interplay between them has rarely been considered (but see Lalonde & Roitberg (2006)). Here, we have provided a minimal model for these dynamics, and developed a basic mathematical account of their consequences.

Recognizing the potential for feedbacks between dormancy and non-equilibrium population dynamics may shed light on a range of biological questions. For example, the apparent rarity of chaos in nature has long intrigued ecologists (Berryman & Millstein, 1989; Hastings *et al*., 1993), although recent analyses suggest that chaotic dynamics are common in certain ecological systems (e.g. plankton and insects), but typically weak or intermittent (Munch *et al*., 2022; Rogers et al., 2022, 2023). We predict that population dynamics with sustained, high-amplitude fluctuations should be very susceptible to invasion by dormancy, with widespread dormancy consequently stabilizing the dynamics. Thus, the evolution of dormancy, which has been achieved in countless taxa through a range of physiological mechanisms, may explain the rarity of these dynamics. Furthermore, our results indicate that, in the absence of other drivers, populations may evolve dormancy strategies that place their dynamics at the edge of stability, consistent with evidence from empirical time-series (Turchin & Ellner, 2000; Rogers *et al*., 2023). Crucially, we show that dormancy is not only stabilizing, but can invade when rare in fluctuating populations, and ESS dormancy strategies can suppress fluctuations while resisting invasion by other strategies. Other mechanisms, such as evolution of lower growth rates (*r*) to suppress chaos, are not similarly evolutionarily stable (for example, higher growth rates can always invade in our model, even if this leads to deterministic fluctuations to extinction) (Parvinen & Dieckmann, 2013). Interestingly, however, the dynamics we identify may be closely related to models of demographic trade-offs, where evolution of high juvenile survival at the expense of lower maturation rates can similarly explain the evolution of stable population dynamics (Ebenman *et al*., 1997).

The interplay of dormancy and fluctuating population dynamics might also provide a mechanism for the coexistence of species or ecotypes with similar – or even identical – niches (e.g. resource use) but contrasting dormancy strategies. Mutual invasibility between high-dormancy strategies that benefit from instability but stabilize dynamics and low dormancy strategies that benefit from stability but destabilize dynamics produces negative frequency-dependence, enabling robust coexistence between different strategies (Fig. 3). Coexistence in our model is possible between species where one falls above the ESS trait value (*α*^⋆^) and the other below (see Fig. 4). This coexistence mechanism can be understood as a “stabilizer-destabilizer trade-off” (Yamamichi & Letten, 2022), a concept recently proposed to generalize the gleaner-opportunist trade-off, which is a fluctuation-dependent coexistence mechanism usually associated with resource competition (Armstrong & McGehee, 1976; Chesson, 1994). The feedback between overcompensatory population dynamics and dormancy produces a stabilizer-destabilizer trade-off without the need to explicitly model fluctuating resource dynamics; thus, we suggest that our framework provides a minimal model of this class of coexistence mechanism.

Finally, we echo Lalonde and Roitberg Lalonde & Roitberg (2006) in speculating that population fluctuations – or the potential of ecological dynamics to fluctuate – may be an underappreciated driver of the evolution of dormancy (see also Verin & Tellier (2018); Ruf & Bieber (2023)). While dormancy is almost always understood as a strategy for avoiding or hedging against harmful abiotic conditions, it is sometimes found in situations with no obvious abiotic driver (Lalonde & Roitberg, 2006; Santangelo *et al*., 2015). Intrinsic population fluctuations might explain these cases, as they create a strong selective advantage for dormancy, but subsequent stabilization of the dynamics would erase signs of the selection pressure.

Our results highlight that a simple, ubiquitous life-history strategy – dormancy – can interact with population and community dynamics to produce complex outcomes. We explored these interactions in a minimal model of a single population, but there is much more to learn about the consequences of dormant states and stages in complex food webs and other multispecies systems. Real-world populations are simultaneously forced by abiotic factors, providing another dimension to be explored. As more ecologists turn their attention to dormancy (Lennon *et al*., 2021; Wisnoski & Lennon, 2021), and return their attention to nonequilibrium population dynamics (Munch *et al*., 2022), there is both opportunity and need to consider how the two interact in natural ecosystems.

## Acknowledgements

We thank Elly Goetz for helpful discussions. This research received support through Schmidt Sciences, LLC.

## Data accessibility

All code used to generate the analysis and figures is available at https://github.com/zacharyrmiller/stabilization_via_dormancy.

## Supplementary Material for

### A Stabilizing effect of dormancy in one-dimensional models

In one-dimensional population models where overcompensation leads to instability, we show that dormancy has a very general stabilizing effect. First, we show that for the class of “separable” models, populations with dormancy are always stable for a wider range of parameters than those without dormancy. Second, we show that the same is true in the limit of small mortality in dormancy (small *m*) regardless of the specific form of the model. Third, we derive the exact bifurcation point for the logistic model with dormancy as a function of *α* and *m*; this stability threshold always increases with increasing dormancy.

#### A.1 Separable models

Many discrete-time models in ecology incorporate density-dependence of the form

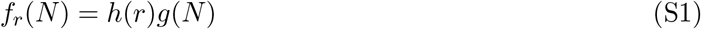

where *h* is an increasing function of the intrinsic growth rate parameter *r >* 0, and *g* is a decreasing function of the population size *N*. We call such models *separable*, because the effect of *r* and *N* can be factored into separate functions. Note that in the main text we define separable models more simply as those where *f*_*r*_(*N*) = *rg*(*N*); the definition in Eq. S1 is slightly more general, but essentially equivalent, as we can always define *ρ* = *h*(*r*) and re-express *h*(*r*)*g*(*N*_*t*_) = *ρg*(*N*_*t*_). For example, the Ricker model is sometimes written as *N*_*t*+1_ = exp *r* 1 − *N*_*t*_*/K N*_*t*_, but using *ρ* = exp(*r*) and rescaling *x* = *rN*, we have *x*_*t*+1_ = *ρ* exp(−*x*_*t*_*/K*)*N*_*t*_. Hereafter, we assume that any separable model is already in the form *f*_*r*_(*N*) = *rg*(*N*).

Consider a separable model with an equilibrium point *N*_*r*_, which is a function of *r*. We assume that there is a unique bifurcation point in *r*, at a value *r*_*c*_, such that *N*_*r*_ is stable for *r < r*_*c*_ and unstable for *r > r*_*c*_. Letting *F*_*r*_(*N*) = *f*_*r*_(*N*)*N*, this implies that

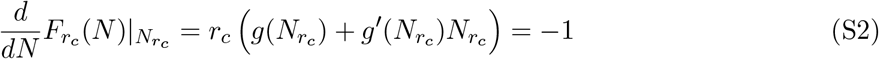

and furthermore 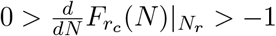 for all *r < r*_*c*_ (Devaney, 2019).

Now we consider the effect of adding dormancy in the model. We consider the family of dynamics *N*_*t*+1_ = *αf*_*r*_(*αN*_*t*_) + (1− *α*)(1− *m*) *N*_*t*_, as in the main text. We assume throughout that *m <* 1. It is convenient to use the change of variable *x* = *αN* and define *F*_*r,α*_(*x*) = *αrg*(*x*) + (1− *α*)(1− *m*) *x*, so that the dynamics can be expressed simply as *x*_*t*+1_ = *F*_*r,α*_(*x*_*t*_).

Let *r*_*c*_(*α*) be the bifurcation point as a function of *α*. We will show that *r*_*c*_(*α*) >*r*_*c*_ for all *α* < 1; that is, populations with dormancy remain stable at higher intrinsic growth rates than those without dormancy. In fact, we establish a quantitative estimate of this stabilizing effect:

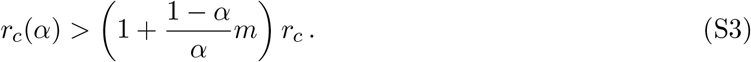

Consider that for a given *α* and *r*, the dynamics with dormancy have an equilibrium *x*_*r,α*_ satisfying *αrg*(*x*_*r,α*_) + (1− *α*)(1− *m*) = 1. For the dynamics without dormancy (*α* = 1), there is a corresponding intrinsic growth rate, which we denote *p*_*r,α*_, such that the dynamics possess the same equilibrium, *x*_*r,α*_. In other words, we define *p*_*r,α*_ to satisfy *p*_*r,α*_ *g*(*x*_*r,α*_) = 1. Combining these two equilibrium conditions, we have that

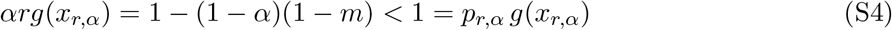

and, in particular, *αr < p*_*r,α*_.

To determine the bifurcation point *r*_*c*_(*α*), we seek the value *r* for which

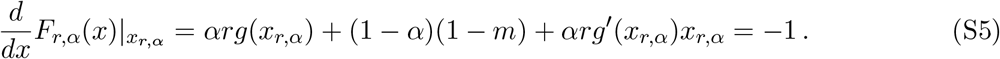

Because *αr* ≤ *p*_*r,α*_ and (1 − *α*)(1 − *m*) > 0, we see that 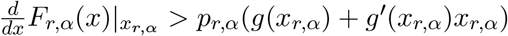. And because we have assumed that 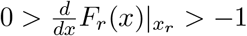 for all *r* < *r*_*c*_, it is clear that *r*_*c*_(*α*) is greater than the value *r* such that *p*_*r,α*_(*g*(*x*_*r,α*_) + *g*^′^(*x*_*r,α*_)*x*_*r,α*_) = − 1. But this last equation is precisely the condition that defines *r*_*c*_ (Eq. S2), so *p*_*r,α*_ = *r*_*c*_. Finally, using Eq. S4, we have that 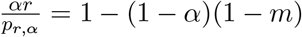 and consequently *p*_*r,α*_ = *r*_*c*_ when 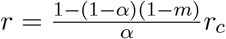. This establishes our quantitative bound, and it follows that *r*_*c*_(*α*) > *r*_*c*_ for all *α* < 1.

#### A.2 Non-separable models

For models where density-dependence is not a separable function of *N* and *r*, we are limited to studying the effect of dormancy on stability for small values of *m*. Specifically, we show that in the limit *m* → 0, dormancy always increases the range of intrinsic growth rates compatible with stability. To see this, we first note that with *m* ≈ 0, the dynamics become

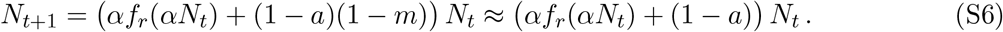

Regardless of the value of *α*, we find that *f*_*r*_(*αN*_*r,α*_) = 1 at equilibrium, implying *αN*_*r,α*_ is equal to a constant, 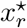, as *α* varies. Now it is straightforward to evaluate the derivative of *F*_*α,r*_(*x*) = *αf*_*r*_(*x*) + (1 − *a*)(1 − *m*) *x* at the equilibrium value 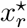:

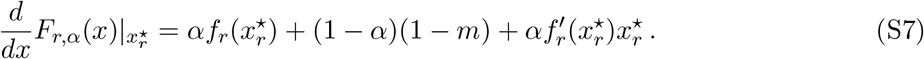

With no dormancy, *r*_*c*_ is defined by the equation 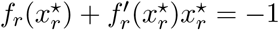. Plugging this condition into Eq. S7, above, gives us 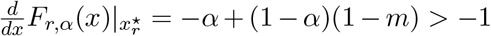, implying that the equilibrium with *α <* 1 is stable at the point where the model without dormancy begins to fluctuate. Thus, the bifurcation point has increased, permitting stability for a greater range of intrinsic growth rates.

#### A.3 Discrete logistic model

For the special case of the discrete logistic model (which assumes separable density dependence), we can exactly solve for the bifurcation point *r*_*c*_(*α*) as function of *α*. As in the previous two cases, we do this by establishing a mapping between the model with parameters *α* and *r* and the model with no dormancy (*α* = 1) and a different intrinsic growth rate.

If we expand the logistic model, as introduced in the main text, and collect linear terms, we can re-express the model as

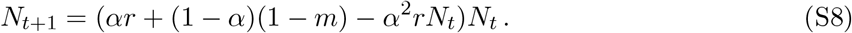

To map these dynamics into the standard logistic model with no dormancy, we introduce the rescaled abundances *x*_*t*_ = *α*^2^*rN*_*t*_. This gives us the dynamics *x*_*t*+1_ = (*αr* + (1 − *α*)(1 − *m*) − *x*_*t*_)*x*_*t*_, and finally defining *r*_eff_ = *αr* + (1 − *α*)(1 − *m*), we see that the dynamics are equivalent to

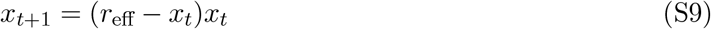

This is a standard logistic model with no dormancy. This transformation only required a re-scaling of the population size; thus, Eq. S8 and Eq. S9 share the same qualitative dynamics.

We can invert the relationship *r*_eff_ = *αr* + (1− *α*)(1−*m*) to solve for the bifurcation point *r*_*c*_(*α*). Using the fact that *r* = 3 is the bifurcation point in the logistic model with no dormancy (Devaney, 2019), we set *r*_eff_ = 3 and solve

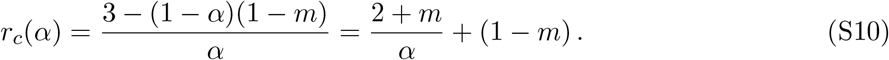

### B Invasion growth rate calculation

As outlined in the main text, we calculate the invasion growth rate (IGR) for a rare ecotype with an active fraction *α*^′^ (“high dormancy”) in a fluctuating resident population with no or lower dormancy (1 ≥ *α* > *α*^′^). First, we must specify the dynamics of the two types. We consider coupling the two populations such that there is no niche differentiation – both types share the same density dependence function, *f*_*r*_, applied to the sum of both active population sizes. This gives us the dynamics

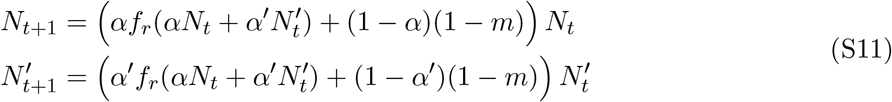

where *N*^′^ denotes the population size of the invader. We treat this population size as nearly zero, so that we can approximate 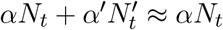. We are interested in calculating the multiplicative growth rate of the invader when the resident population size, *N*_1_, *N*_2_, …, has reached a stationary state (limit cycle or chaotic attractor). At a stationary state, the resident population is neither growing nor shrinking in the long term; thus, the per capita growth rate of the resident has a geometric mean of one over the long term. Equivalently, defining *h*_*t*_ = *αf*_*r*_(*αN*_*t*_) + (1 − *α*)(1 − *m*), we have that 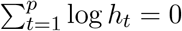, where *p* is the period of the resident population fluctuations (for chaotic dynamics, we take *p* → ∞).

The geometric mean (per generation) growth rate of the invader in this context is straightforwardly given by

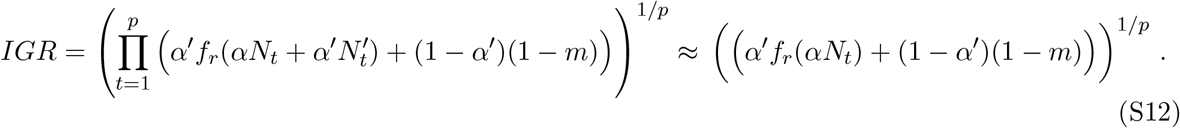

To manipulate the *IGR*, it is more convenient to work with its logarithm. We have

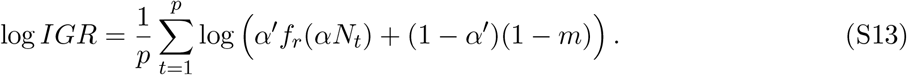

With some algebra, we can express the log IGR in terms of the resident growth rate, *h*_*t*_:

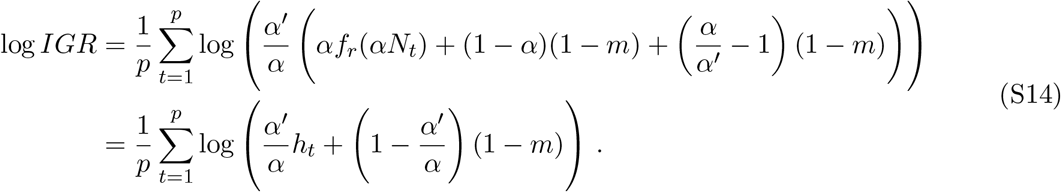

We can now apply Jensen’s inequality (Ruel *et al*., 1999) term-by-term to the convex function − log(*x*), to obtain the lower bound

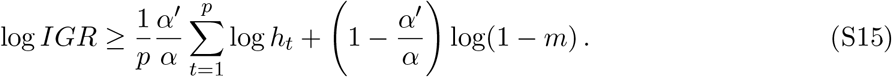

By assumption, the first term 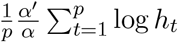 is zero. The second term is negative for *m >* 0; we deem this the mortality cost of dormancy.

The difference between the left- and right-hand sides of Eq. S15 is often called the “Jensen gap” (Liao & Berg, 2019). We denote this quantity *J* and refer to it as the bet-hedging benefit of dormancy. *J* is zero if and only if *h*_*t*_ is constant for all *t*. It is easy to see that if 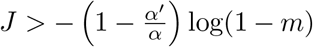, then log *IGR >* 0 and the invader population will grow. Whether this is the case depends on the distribution of *h*_*t*_, the value of *m*, and the relative values of *α* and *α*^′^. However, we can prove that there is a always a choice of *m* sufficiently small so that log *IGR >* 0, for any fixed *α*^′^ *< α* and any non-constant *h*_*t*_. From Liao & Berg (2019), we have the lower bound

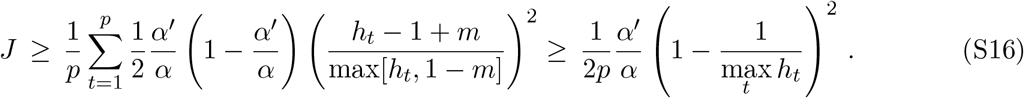

The second inequality in Eq. S16 provides a lower bound that has no dependence on *m*. Note that this bound simply assumes the existence of some *h*_*t*_ > 1 > 1 − *m*, which is guaranteed by our assumptions. Thus, as *m* → 0, we see that the mortality cost also shrinks to zero, while *J* remains greater than a nonzero constant. We have made no attempt to optimize the bound in Eq. S16; however, this simple lower bound is sufficient to show that log *IGR >* 0 for *m* sufficiently small.

To explore the robustness of the IGR advantage of dormancy in the presence of a mortality cost, we numerically calculated the IGR for an invader in the logistic model with varying *m* and *α*^′^. For simplicity, we assumed no dormancy in the resident population. Results for four values of *r* are shown in Fig. S1. We observe that positive log IGRs (successful invasion) are achieved for non-negligible values of *m*. As *r* increases, fluctuations in the resident population increase in amplitude, conferring a large IGR to dormancy strategies, even when mortality in dormancy, *m*, is considerable.

**Figure S1:**
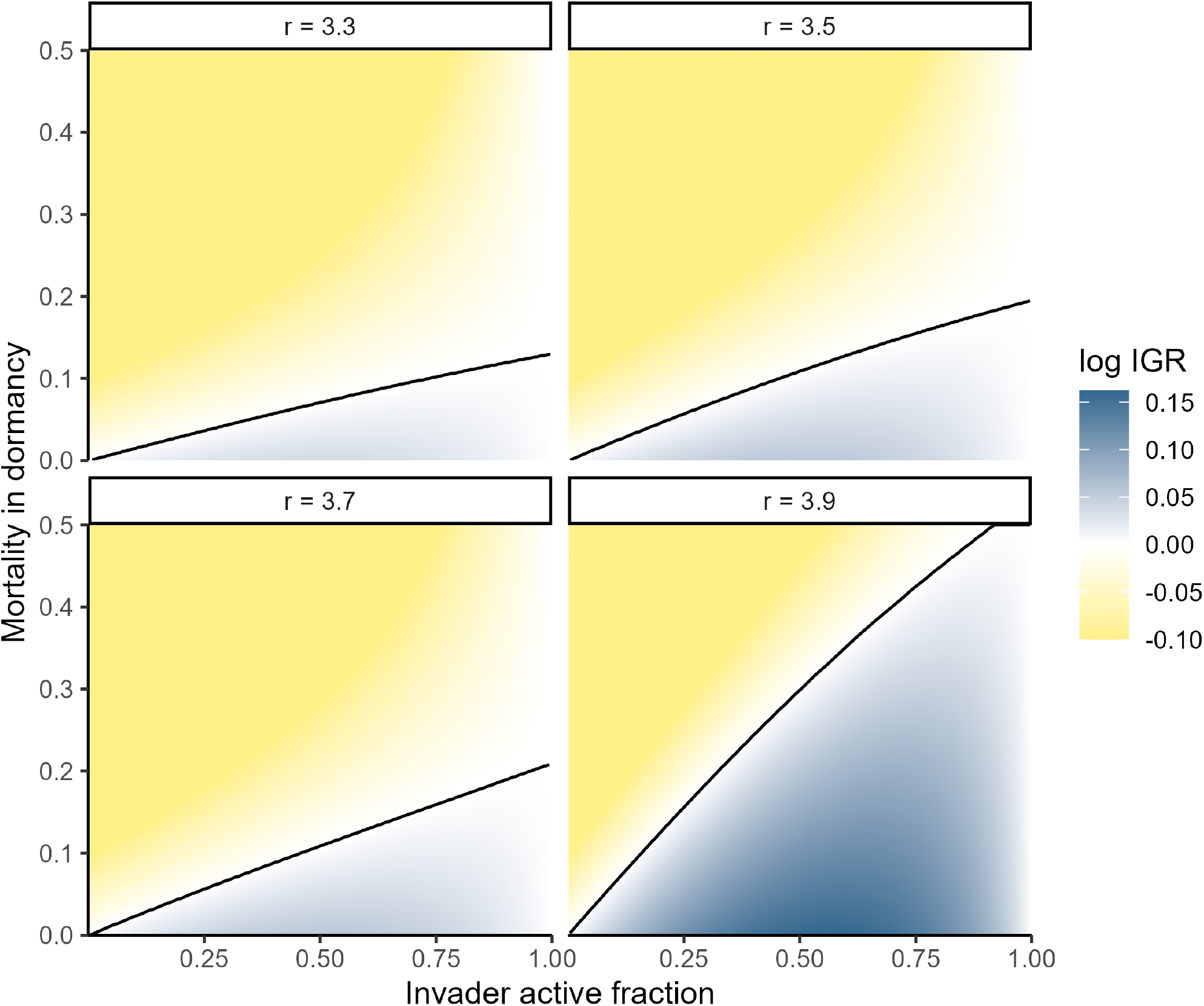
Invasion growth rate as a function of invader active fraction, *α*^′^ (x axis) and mortality in dormancy, *m* (y axis). The resident population has no dormancy (*α* = 1) and is taken to be at a stationary state (i.e. its dynamical attractor). Results for different values of *r*, shared by both resident and invader, are shown in each panel. Black line indicates the break between positive and negative log IGR. Note that log IGR values below −0.1 (corresponding to ≈ 10% per generation population decline) were truncated for clearer visualization.

### C Invasion growth rates in the continuous-time model

As discussed in the main text (Box 1), we can also consider the interaction between dormancy and fluctuating population dynamics in continuous time. In this setting, species interactions are required to produce fluctuating dynamics; limit cycles are only possible with at least two interacting species, and chaos requires at least three.

Here, we show that dormancy also confers a growth rate advantage when population dynamics fluctuate in this alternative setting.

Consider some very generic community dynamics

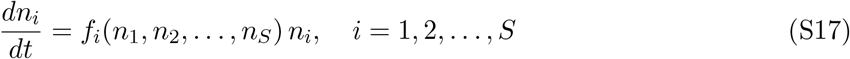

for *S* interacting species. We assume that the resident community has reached some dynamical attractor where the time-averaged growth rate for each species approaches zero:

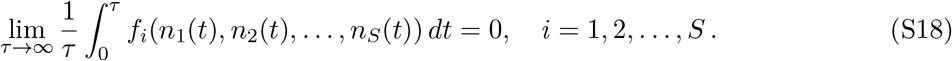

Now we consider an invader that is identical to some resident species *j*, but transitions in and out of a dormant state. More precisely, the dynamics of the resident community and the invader are given by

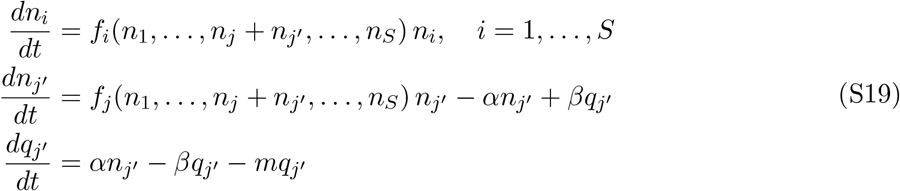

where 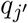 is the dormant population size of the invader, and *α, β >* 0 are the rates at which the invader species transitions in and out of the dormant state, respectively.

To evaluate the invasion growth rate of the invader with dormancy, we again suppose that 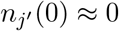 (and 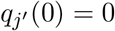). We will also assume that *m* ≈ 0 for simplicity, although, as in the discrete-time case, substantial mortality risk in dormancy may ultimately outweigh the benefits of dormancy.

With these assumptions, we are interested in the dynamics of

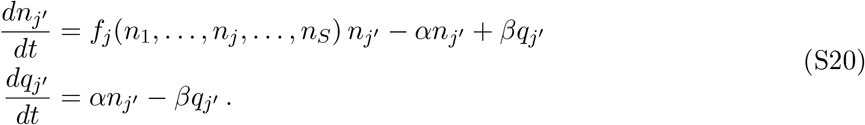

Because the effect of the invader on the resident dynamics is initially negligible, we can simply treat *f*_*j*_(*n*_1_, *n*_2_, …, *n*_*j*_, …, *n*_*S*_) = *f* (*t*) as an external forcing characterized by

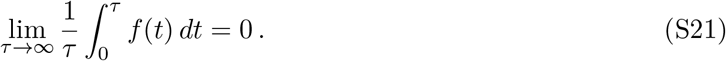

While we are unable to characterize these dynamics in full generality, we show here that the invader population always grows in two limits: *α, β* → ∞, corresponding to very frequent transitions in and out of dormancy, and *α, β* → ∞, corresponding to very infrequent transitions (in both cases, we assume the ratio *α/β* approaches a non-zero constant). For convenience, we refer to these cases as “fast” and “slow” dormancy dynamics, respectively. These two cases bookend all possible scenarios. Based on the consistent behavior between these two extremes, we conjecture that dormancy invades for all choices of *α* and *β*.

#### C.1 Fast dormancy dynamics

To characterize the IGR in the limit of fast dormancy dynamics, it is convenient to define the new variables 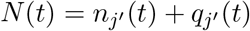, the total (active and dormant) population size of the invader, and 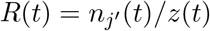, the fraction of the population that is active at time *t*. After some algebra, we find the dynamics

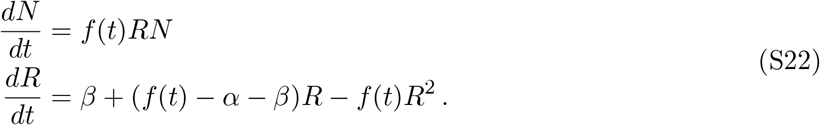

Eq. S22 makes clear that the fate of the invader depends upon the sign of 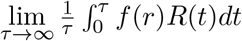. If this quantity is positive, the invader population will grow in the long run.

Under the assumption of fast dormancy dynamics, Eq. S22 can be treated as a “fast-slow” system (Klonowski, 1983). We make this more explicit by defining *ϵ* = 1*/β* and *γ* = *α/β*. Then we have

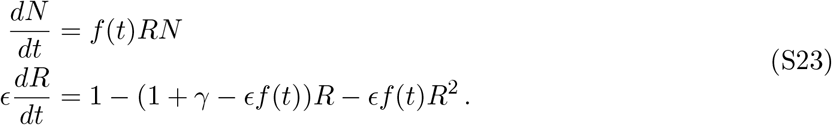

We can obtain a reduced system in the limit *ϵ* → 0, by taking *R*(*t*) to be at its (stable) equilibrium as a function of *f* (*t*) (Klonowski, 1983). The dynamics for *R*(*t*) have the unique stable equilibrium point

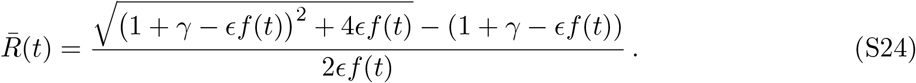

Using this reduction, we find that the total population dynamics are approximated by

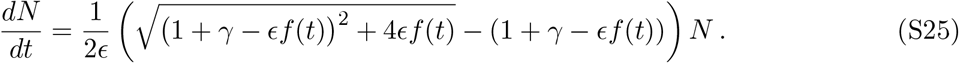

It is straightforward to verify that 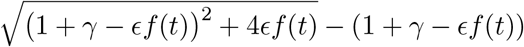 is a convex function of *f* (*t*) (e.g., by computation of the second derivative). Thus, we can once again use Jensen’s inequality to establish that

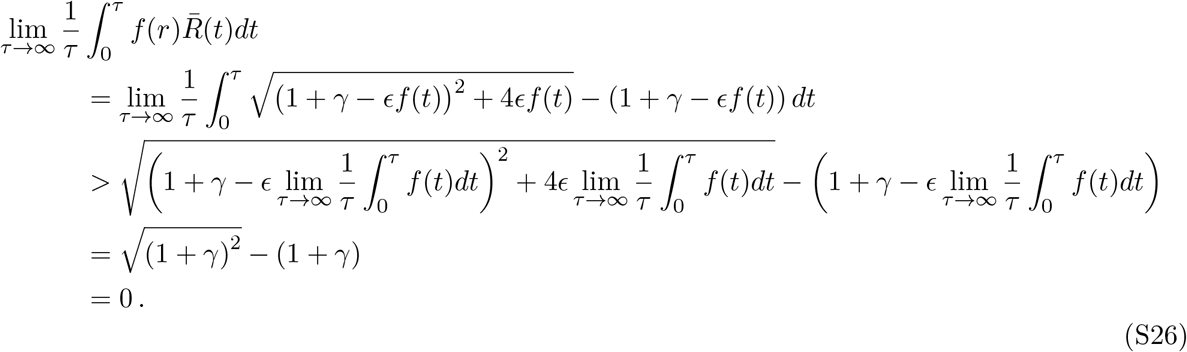

As in the discrete-time case, we find that non-linear averaging over times of population growth and decline confers a benefit to dormancy. It is interesting to note that even a stochastic dormancy strategy – in which individuals go dormant and become active with a constant probability per time, with no cues from the environment – is sufficient to generate a positive correlation between *R*(*t*), the fraction of the population that is active, and *f* (*t*), the growth rate of the active population. This is essentially what is expressed by 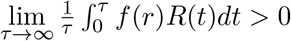. We note that more active dormancy strategies involving sensory mechanisms would be expected to strengthen this correlation if they are adaptive. Thus, more active forms of dormancy should exhibit even larger invasion growth rates.

#### C.2 Slow dormancy dynamics

As another extreme case, we can consider the limit *α, β* → 0. As in the last section, we define *γ* = *α/β*, and now we let *ϵ* = *β*. To treat the dynamics in this case, it is convenient to introduce matrix notation: we define 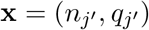 and

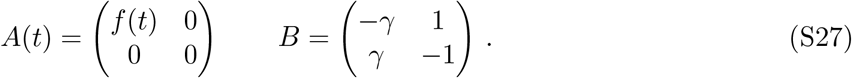

With these conventions, we can re-write Eq. S20 as

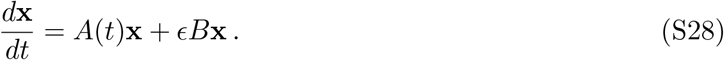

We can view these dynamics as the (linear) dynamics of the resident species plus a small perturbation. By transforming this system into Lagrange standard form, we can apply the method of averaging to obtain an approximation of the dynamics (Verhulst, 2012). Intuitively, because the flows in and out of the dormant state are slow, the long-term dynamics of the invader population can be treated as though they depend on an appropriate average of the community fluctuations.

First, we obtain the fundamental matrix for the unperturbed system, 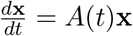, which takes a very simple form:

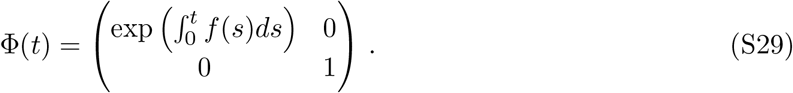

Next we introduce the comoving coordinates **y** = Φ^−1^(*t*)**x**. The dynamics of **y** are found to be

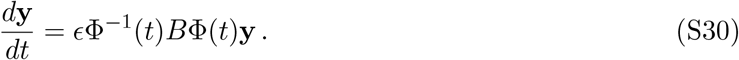

Eq. S30 is now in a form where we can apply time-averaging. The long-term dynamics of Eq. S30 can be approximated by

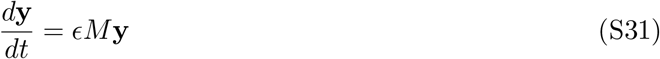

Where

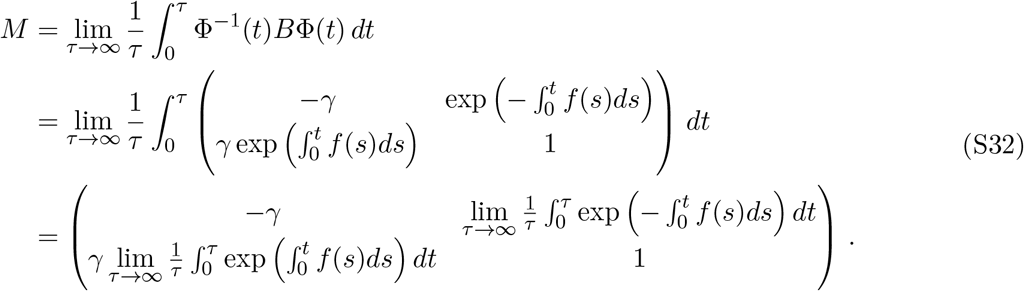

In order to evaluate the integrals in *M*, we consider our assumption that the resident species *j* is fluctuating in a stationary fashion. This implies that the time-average of *n*_*j*_ converges to a constant:

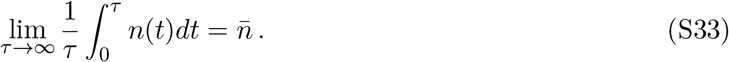

But we can express 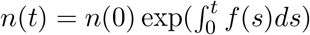, so, equivalently,

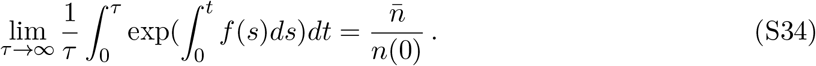

We define 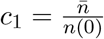. Eq. S34 gives us one of the integrals in *M*. For the other integral, we recognize that *z*^−1^ is a convex function. Yet again, we can apply Jensen’s inequality to obtain

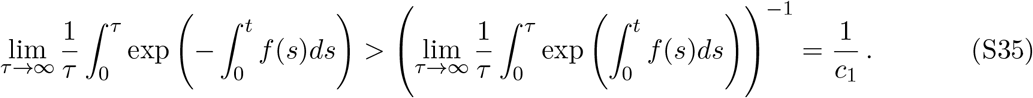

Denoting the value of this integral as *c*_2_ > 1*/c*_1_, we finally obtain

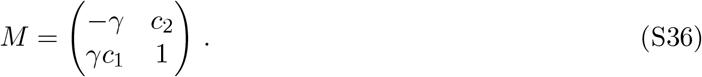

Up to this point, we have established that the dynamics of the invader, written in the coordinates **y**, can be approximated by the matrix differential equation 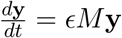. The solutions depend on the eigenvalues of *M* :

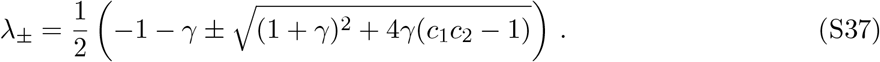

Because we have that *c*_2_ > 1*/c*_1_, we must have *c*_1_*c*_2_ > 1, and therefore the eigenvalue *λ*_+_ is positive. This eigenvalue corresponds to the eigenvector

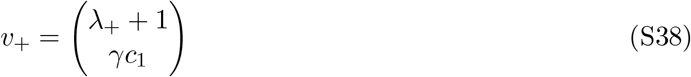

which therefore grows exponentially. Our initial condition 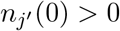 and 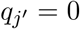 necessarily has a positive projection on this eigenvector. Thus, 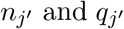 grow in the long term, and the invader has a positive IGR.

### D Alternative model: the Ricker model

In this section, we illustrate that results shown in the main text for the logistic model apply to another well-known population model, the Ricker model (Ricker, 1954; Geritz & Kisdi, 2004). We define the Ricker model dynamics with dormancy as

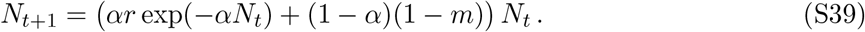

Like the logistic model, the basic Ricker model is overcompensating, and the model with no dormancy exhibits a bifurcation at *r* = exp(2), followed by a period-doubling cascade and the onset of chaotic dynamics for larger values of *r*. Unlike the logistic model, the Ricker model assumes no maximum population size. The Ricker model is often considered to be more biologically plausible and is commonly used in fisheries and other applications (Subbey *et al*., 2014).

Here, we reproduce for the Ricker model the bifurcation diagram (Fig. S2) and pairwise-invasibility plot (Fig. S3), shown in the main text for the logistic model (Figs. 2 and 4). These plots are qualitatively identical to those shown for the logistic model, illustrating the generality of our results.

**Figure S2:**
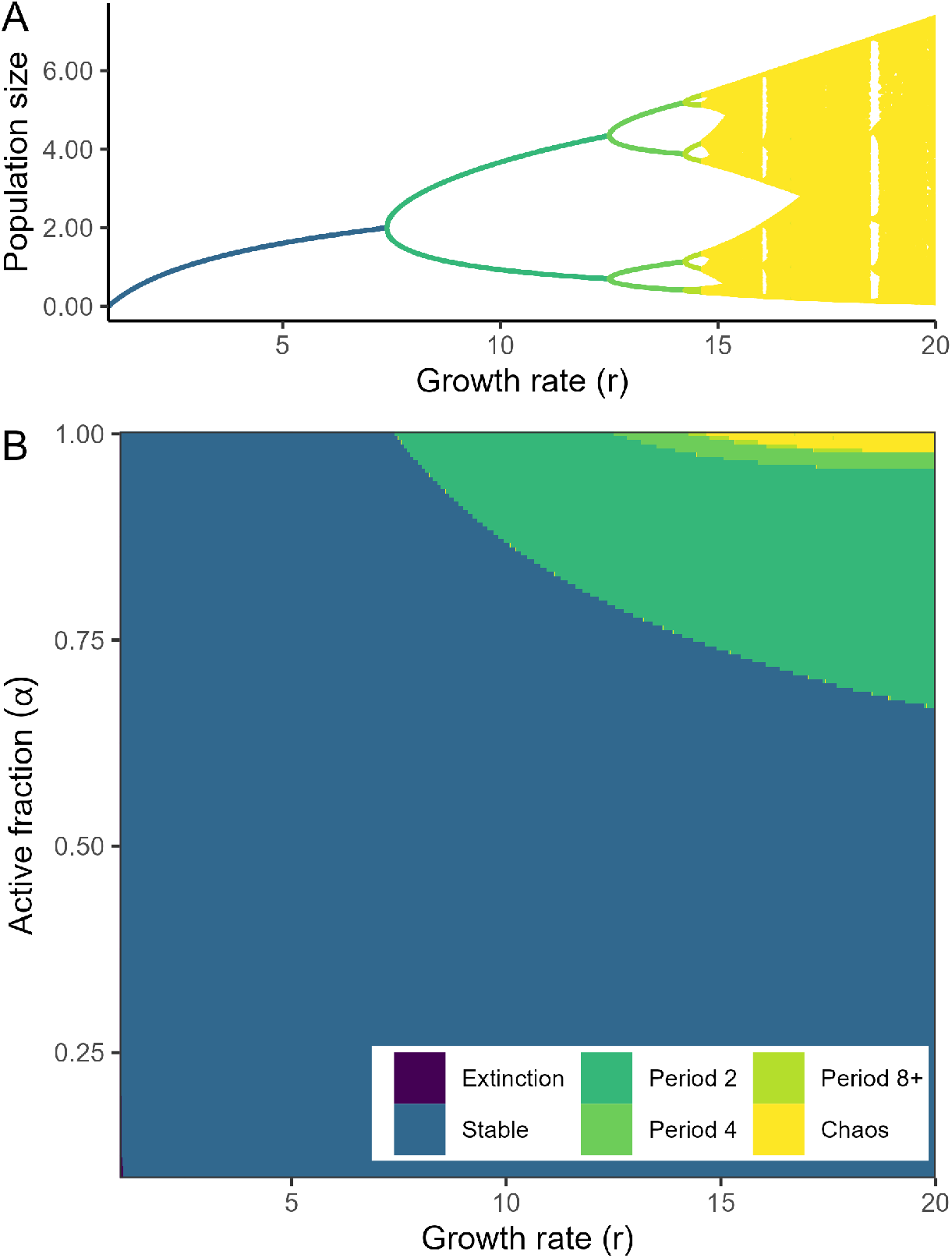
Bifurcation diagrams for the Ricker model (Eq. S39). (A) Steady state population size(s) are plotted against *r* for the standard Ricker model with *α* = 1. For *r <* exp(2) the model possesses a stable equilibrium. At *r* = exp(2) the dynamics undergo a bifurcation and stable oscillations (green) emerge. Furthering increasing *r* leads to the onset of chaos. (B) Qualitative steady state dynamics for different combinations of *r* and *α* (with *m* = 0.01). Dormancy increases from top to bottom. We determined the dynamics for each parameter combination by directly iterating Eq. S39. For lower values of *α* (more dormancy), population fluctuations arise at higher growth rates (*r*), indicating a stabilizing effect of dormancy.

**Figure S3:**
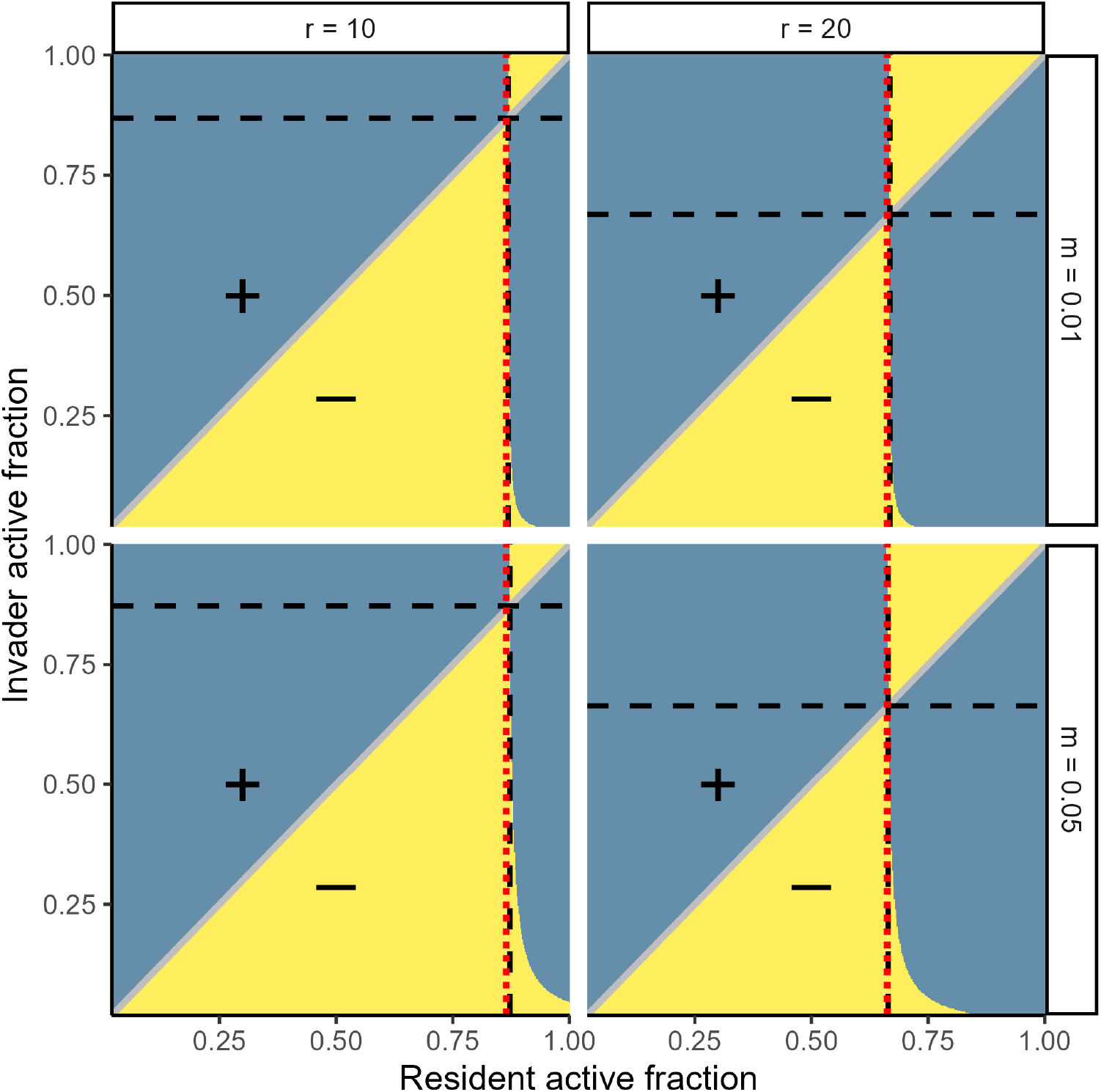
Pairwise invasibility plots for the trait value *α* in the Ricker model (Eq. S39) with different fixed values of *r* and *m*. Blue regions indicate that the active fraction *α*^′^ on the y-axis can invade the resident active fraction *α* on the x-axis. Yellow regions indicate that the resident can resist invasion. For each combination of *r* and *m*, there is a unique value *α*^⋆^ that is not invasible by any other *α* value (vertical dashed line), and that can invade every other value (horizontal dashed line). These are evolutionary stable strategies (ESS). As discussed in the main text, the ESS typically sits close to – but just above – the value *α*_*c*_ where the dynamics begin to fluctuate (red dotted lines). As for the logistic model (Fig. 4, main text) we observe that *α*^⋆^ ≈ *α*_*c*_; in fact, the correspondence is even closer in this case.

## Notes

https://github.com/zacharyrmiller/stabilization_via_dormancy

### Competing Interest Statement

The authors have declared no competing interest.

